# High affinity binding of SARS-CoV-2 spike protein enhances ACE2 carboxypeptidase activity

**DOI:** 10.1101/2020.07.01.182659

**Authors:** Jinghua Lu, Peter D. Sun

## Abstract

A novel coronavirus (SARS-CoV-2) has emerged to a global pandemic and caused significant damages to public health. Human angiotensin-converting enzyme 2(ACE2) was identified as the entry receptor for SARS-CoV-2. As a carboxypeptidase, ACE2 cleaves many biological substrates besides Ang II to control vasodilatation and permeability. Given the nanomolar high affinity between ACE2 and SARS-CoV-2 spike protein, we wonder how this interaction would affect the enzymatic activity of ACE2. Surprisingly, SARS-CoV-2 trimeric spike protein increased ACE2 proteolytic activity ~3-10 fold when fluorogenic caspase-1 substrate and Bradykinin-analog peptides were used to characterize ACE2 activity. In addition, the enhancement was mediated by ACE2 binding of RBD domain of SARS-CoV-2 spike. These results highlighted the altered activity of ACE2 during SARS-CoV-2 infection and would shed new lights on the pathogenesis of COVID-19 and its complications for better treatments.

## Introduction

The novel coronavirus SARS-CoV-2 has emerged as an unprecedented global pandemic ^1, 2, 3^ and as of June 8, 2020, there are nearly 7 million confirmed cases worldwide, leading to 400,857 deaths(WHO). The infection of SARS-CoV-2 causes fever, dry cough, severe respiratory illness and pneumonia, a disease recently named COVID-19^4^. Pathological studies have revealed all features of diffuse alveolar damage (DAD) with excessive fluid in the lungs and critically ill patients often need ventilators to support oxygen. Furthermore, a frequent complication of COVID-19 that blood clotted abnormally is observed in many hospitalized patients^5^. Therefore, there is an urgent need for a mechanistic understanding of the pathogenicity of SARS-CoV-2 and its concomitant complications to be able to treat hospitalized patients.

During the sudden emergency and rapid spread of SARS-CoV-2, global efforts have identified human angiotensin-converting enzyme 2(ACE2) as the entry receptor for this new coronavirus^2^, which was also the entry receptor for SARS-CoV^6, 7, 8^. Structural studies revealed that SARS-CoV-2 spike (S) glycoprotein binds ACE2 with higher affinity (~15-40nM) than SARS-CoV spike protein^9, 10, 11, 12^. The overall structure of SARS-CoV-2 S resembles that of SARS-CoV S with the RBD domain of one protomer in spike protein trimer tightly interacting ACE2 extracellular enzymatic domain. Physiologically, ACE2 is a zinc metalloprotease (carboxypeptidase), a homolog to dipeptidase angiotensin-converting enzyme (ACE) but with different substrate specificity ^13^. ACE cleaves the C-terminal dipeptide from Ang I to produce the potent vasopressor octapeptide Ang II, whereas ACE2 further cleaves the carboxyl-terminus to produce Ang 1-7 to deactivate Ang II. Therefore, ACE and ACE2 together play important roles in regulating vasoconstriction and vasodilatation in the rennin-angiotensin system (RAS). In the meanwhile, ACE and ACE2 plays critical role in kinin-kallikrein system to control vascular permeability and vasodilatation^14^. ACE deactivates Bradykinin nonapeptide, the ligand for constitutively expressed bradykinin receptor B2. Bradykinin can be further processed by carboxypeptidase N or M to form des-Arg9-bradykinin, which is a potent ligand for inflammation-inducible bradykinin receptor B1^15^. Beyond Renin-angiotensin and kinin-kallikrein systems, ACE2 also cleaves other biological peptides such as Apelin-13 that activates apelin receptor to cause vasodilatation. However, there is limited information on the level of these substrates during SARS-CoV-2 infection given the fact that ACE2 is the major entry receptor for this new coronavirus.

Compared to relatively low expression of ACE, ACE2 is predominantly expressed in the lung on pneumocytes II, which explains that lung is the major organ infected by SARS-CoV-2. Critical clinical observations showed that COVID-19 patients often had dyspnea and accumulation of fluid in the lungs representing local angioedema, which clearly suggests the pathological involvement of vascular permeability and vasodilatation during SARS-CoV-2 infection. However, there was no direct assessment of ACE2 enzymatic activity during coronavirus infection. Here we examined how the binding of SARS-CoV-2 spike protein would affect the intrinsic enzymatic activity of ACE2 using one well characterized fluorogenic ACE2 substrate, the caspase-1 substrate (Mca-YVADAPK-Dnp)^13^. In comparison, ACE substrate, a bradykinin analog (Mca-RPPGFSAFK-Dnp) was also included in the enzymatic activity assessment^16^. To our surprise, SARS-CoV-2 spike enhanced ACE2 proteolytic activity to cleave Mca-YVADAPK-Dnp, which was mediated by ACE2 binding of RBD. Furthermore, SARS-CoV-2 RBD enhanced ACE2 activity to hydrolyze the bradykinin-analog better than SARS-CoV RBD. Measurements of kinetic constants also showed that SARS-CoV-2 spike protein altered the binding affinity (*K*_m_) of ACE2 to the caspase-1 substrate or Bradykinin-analog. We propose that this new line of evidence that SARS-CoV-2 spike protein significantly alters ACE2 activity and specificity could be clinically relevant to the understanding of the pathogenesis of COVID-19.

## Results

### SARS-CoV-2 spike protein enhances ACE2 activity

Genetic analysis revealed that SARS-CoV-2 was highly homologous to severe acute respiratory syndrome coronavirus (SARS-CoV) that infected human cells with angiotensin-converting enzyme 2(ACE2) receptor^2, 8^. After the outbreak of COVID-19, ACE2 was quickly identified as the entry receptor for SAR-Cov-2 as well. Further structural studies demonstrated that both SARS-CoV-2 and SARS-CoV utilized their RBD domain to interact with ACE2 at a very similar binding mode. However, SARS-CoV-2 spike protein displays a much higher binding affinity to ACE2 compared with SARS-CoV. Given the nanomolar affinity between ACE2 and SARS-CoV-2 S spike protein, we wonder how the binding of SARS-CoV-2 spike protein to ACE2 would affect its enzymatic activity since ACE2 is a critical component in both the Rennin-angiotensin and Kinin-kallikrein system as a carboxypeptidase^14, 17^.

As previously reported, ACE2 efficiently hydrolyzes biological peptides with a consensus sequence of Pro-X(1-3 residues)-Pro-hydrophobic at P5-P1’ positions^13^, with cleavage between proline and the hydrophobic residue as exemplified by Ang II (DRVYIHP↓F) and des-Arg^9^-BK (RPPGFSP↓F). ACE2 also cleaves peptides with a basic residue at P1’ position such as Dynorphin A (YGGFLRRIRPKL↓K) and Neurotensin 1-8(pE-LYENKP↓R). To rapidly and continuously assess enzyme activity, fluorogenic peptide substrates were normally used. Therefore, we monitored ACE2 proteolytic activity by measuring the degradation of fluorogenic capase-1 substrate Mca-YVADAPK(Dnp)^13^. In addition, a bradykinin derivative Mca-RPPGFSAFK(Dnp)-OH^16^ that was developed as endothelin-Converting enzyme-1 (ECE1) and ACE substrate was also analyzed.. Indeed, ACE2 efficiently hydrolyzed Mca-YVADAPK(Dnp) but cleaved bradykinin analog Mca-RPPGFSAFK(Dnp)-OH less (**Figure 1 A and B**). Within 2 hours, the cleavage of Mca-YVADAPK (Dnp) by ACE2 produced 20,000 relative fluorescence units (RFU), whereas the cleavage of Mca-RPPGFSAFK(Dnp)-OH by ACE2 produced 2000 RFU. Surprisingly, the addition of SARS-CoV-2 spike protein at 14ug/ml concentration to the enzymatic assays significantly enhanced ACE2 proteolytic activity. Within 2 hours, the hydrolysis of Mca-YVADAPK(Dnp) by ACE2 produced ~70,000 RFU in the presence of SARS-CoV-2 spike protein (**Figure 1A**). Similarly, SARS-CoV-2 spike protein enhanced ACE2 cleavage of bradykinin analog Mca-RPPGFSAFK(Dnp)-OH to produce ~3 times more RFU within 2 hours (**Figure 1B**). Previous enzymatic assays showed that ACE2 activity increased with the concentrations of NaCl between 0.15M to 1M. We then carried out the same enzymatic assays at increased concentration of NaCl. Consistently, SARS-CoV-2 spike protein enhanced ACE2 activity at 0.3M and 1M NaCl and enabled ACE2 to cleave bradykinin-analog as well (**supplementary Figure 1**), indicating the enhancement was resulted by the interaction between SARS-CoV-2 spike protein and ACE2.

**Figure 1.**
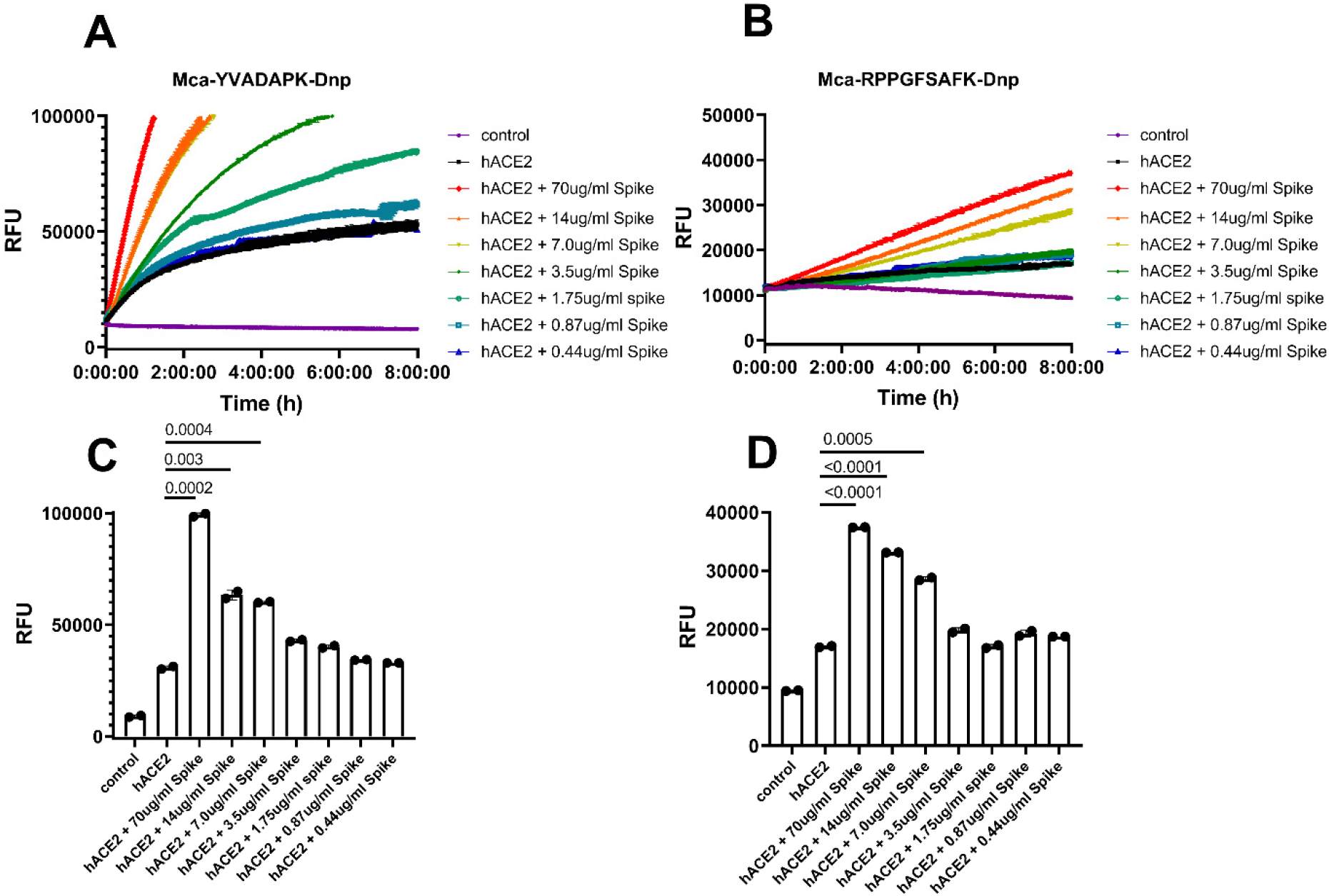
SARS-CoV-2 spike protein enhanced ACE2 cleavage of fluorogenic caspase-1 substrate and bradykinin analog in a concentration dependent manner. A) and B) Kinetic reading of relative fluorescence units (RFU) monitoring the hydrolysis of Mca-YVADAPK-Dnp(20uM) and Mca-RPPGFSAFK-Dnp (20uM) in the presence of SARS-CoV-2 spike protein at indicated final concentrations in the enzymatic assays. The maximum reading is limited to 100000 RFU on Synergy H1 plate reader. C) and D) Comparison of RFU generation during the cleavage of Mca-YVADAPK-Dnp and Mca-RPPGFSAFK-Dnp at 1.5h and 8 h, respectively.

Furthermore, the enhancement of ACE2 activity by SARS-CoV-2 spike protein was at a concentration dependent manner. In the enzymatic assays, we added various concentrations of SARS-CoV-2 spike protein ranging from 70ug/ml to 0.4ug/ml. The dilution of SARS-CoV-2 spike protein in the assays gradually mitigated the enhancement of ACE2 activity with a clear transition between 3.5ug/ml and 7ug/ml of SARS-CoV-2 spike protein (**Figure 1 C and D**). The dose dependent enhancement of ACE2 activity by SARS-CoV-2 spike protein was in line with the binding affinity between ACE2 and SARS-CoV-2 spike protein, which was reported to be 14.7nM, equivalent to ~7ug/ml. Previous studies showed that SARS-CoV infection may lead to the internalization of ACE2 receptor and thus decrease the cell surface level of ACE2^6^. However, examining the lung tissue 3 days post infection of SARS-CoV-2 virus in ACE2 transgenic mice showed colocalization of SARS-Cov-2 spike protein and ACE2 receptor on the cell surface^7, 18^, further supporting SARS-CoV-2 spike formed a stable interaction with ACE2 for the cell entry. In addition, the lasting interaction between SARS-CoV-2 spike and ACE2 on the cell surface suggests that the observed enhancement of ACE2 activity in solution might happen *in vivo* as well.

### SARS-CoV-2 RBD enhanced ACE2 activity better than SARS-CoV RBD

SARS-CoV-2 and SARS-CoV spike protein were highly homologous in their prefusion trimeric architectures with one receptor binding domain (RBD) for ACE2 binding. However, SARS-CoV-2 spike protein binds to ACE2 at ~5-10 fold higher affinity than SARS-CoV spike^9, 11^. We then determined whether RBD domain itself would be sufficient to boost ACE2 activity. The results showed that both SARS-CoV-2 RBD and SARS-CoV RBD enhanced ACE2 cleavage of caspase-1 substrate(**Figure 2 A and B**), demonstrating that SARS-CoV-2 and SARS-CoV RBD alone is sufficient to enhance ACE2 activityACE2. The enhancement of ACE2 activity by SARS-CoV-2 RBD protein showed a concentration dependent saturation with half maximal enhancement at ~70nM, whereas SARS-CoV RBD protein enhances ACE2 activity almost linearly between 63nM and 1000nM with half maximal enhancement at ~170nM (**Figure 2E**). Nevertheless, only SARS-CoV-2 RBD but not SARS-CoV RBD could increase the cleavage of Mca-RPPGFSAFK(Dnp)-OH by ACE2(**Figure 2 C, D and F**). As recently reported, SARS-CoV-2 RBD bound to ACE2 at nanomolar affinities(~30-50nM), while SARS-CoV RBD bound to ACE2 at sub-micromolar affinities(~180-400nM). The different capabilities of SARS-CoV-2 and SARS-CoV RBD proteins to enhance ACE2 enzymatic activity suggested that a more stable interaction between RBD and ACE2 was necessary for its cleavage of non-optimal substrates such as bradykinin analog.

**Figure 2.**
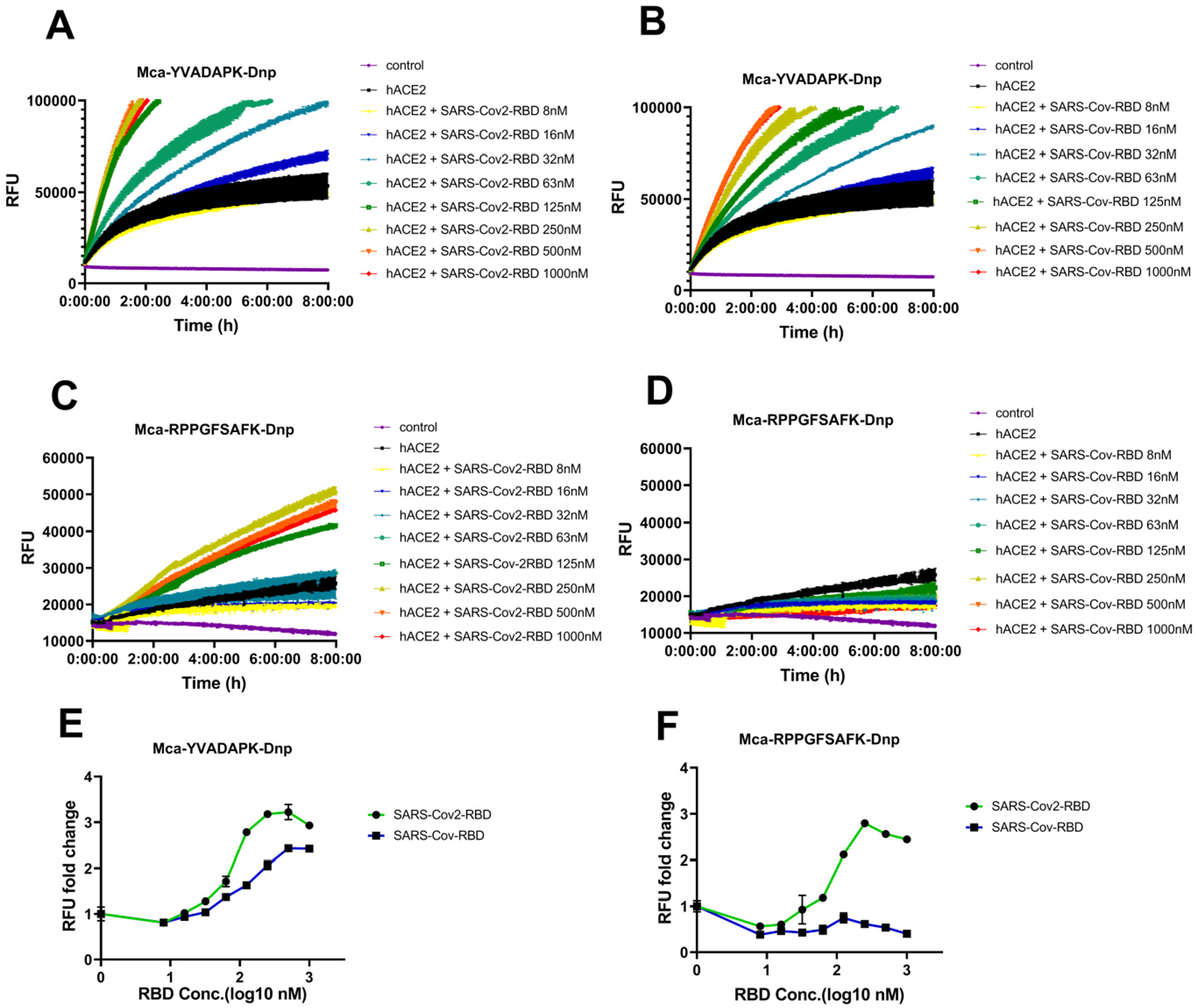
SARS-CoV-2 RBD but not SARS-CoV RBD enabled ACE2 to cleave bradykinin analog. A) and B) hydrolysis of Mca-YVADAPK-Dnp(20uM) in the presence of SARS-CoV-2 RBD and SARS-CoV RBD proteins at indicated concentrations. C) and D) hydrolysis of Mca-RPPGFSAFK-Dnp (20uM) in the presence of SARS-CoV-2 RBD and SARS-CoV RBD proteins at indicated concentrations. E) Fold change of RFU during Mca-YVADAPK-Dnp cleavage in the presence of SARS-CoV-2 RBD or SARS-CoV RBD at the time point of 1.5h due to instrument overflow. F) Fold change of RFU during Mca-RPPGFSAFK-Dnp cleavage in the presence of SARS-CoV-2 RBD or SARS-CoV RBD at time point of 8h. All RFU readings at different concentrations of RBD proteins were normalized to that of ACE2 cleavage without RBD proteins correspondingly.

### Determination of kinetic constants of ACE2 in the presence of SARS-CoV-2 spike

To further characterize how efficiently ACE2 cleaves fluorogenic caspase-1 substrate and bradykinin analog in the presence of SARS-CoV-2 spike protein, we determined the kinetic constants for ACE2, particularly under the physiological condition at pH7.5 with 150mM NaCl. The measurements were carried out with 20ng ACE2 in the presence or absence of 14ug/ml SARS-CoV-2 spike protein and the hydrolysis was limited to 15% product formed as initial velocity conditions. Michaelis-Menten plots showed that SARS-CoV-2 spike protein resulted in a ~4 fold reduction of *K*_m_ from 10.2uM to 2.4uM for ACE2 hydrolysis of Mca-RPPGFSAFK(Dnp)-OH(**Figure 3 and Table1**). In the meanwhile, SARS-CoV-2 spike protein mildly reduced the *K*_m_ for ACE2 hydrolysis of Mca-YVADAPK(Dnp) from 46.6uM to 28.2uM. More dramatically, *k*_cat_/*K*_m_ value for ACE2 hydrolysis of Mca-RPPGFSAFK(Dnp)-OH was increased ~10 times in the presence of SARS-CoV-2 spike protein. As a comparison, *k*_cat_/*K*_m_ value for ACE2 hydrolysis of Mca-YVADAPK(Dnp) was boosted moderately to 3 fold. Furthermore, ACE2 showed higher activities at 0.3M NaCl as previously reported. It is worth mention that *k*_cat_/*K*_m_ value for ACE2 hydrolysis of bradykinin analog Mca-RPPGFSAFK(Dnp)-OH in the presence of SARS-CoV-2 spike protein was 7.55 x 10^3^ M^−1^s^−1^, which is slightly higher than that of ACE2 hydrolysis of Angiotensin I decapeptide (DRVYIHPFH↓L) that was a biological substrate for ACE2^13^. Taken together, SARS-CoV-2 increased the substrate binding affinities and catalytic rate of ACE2.

**Figure 3.**
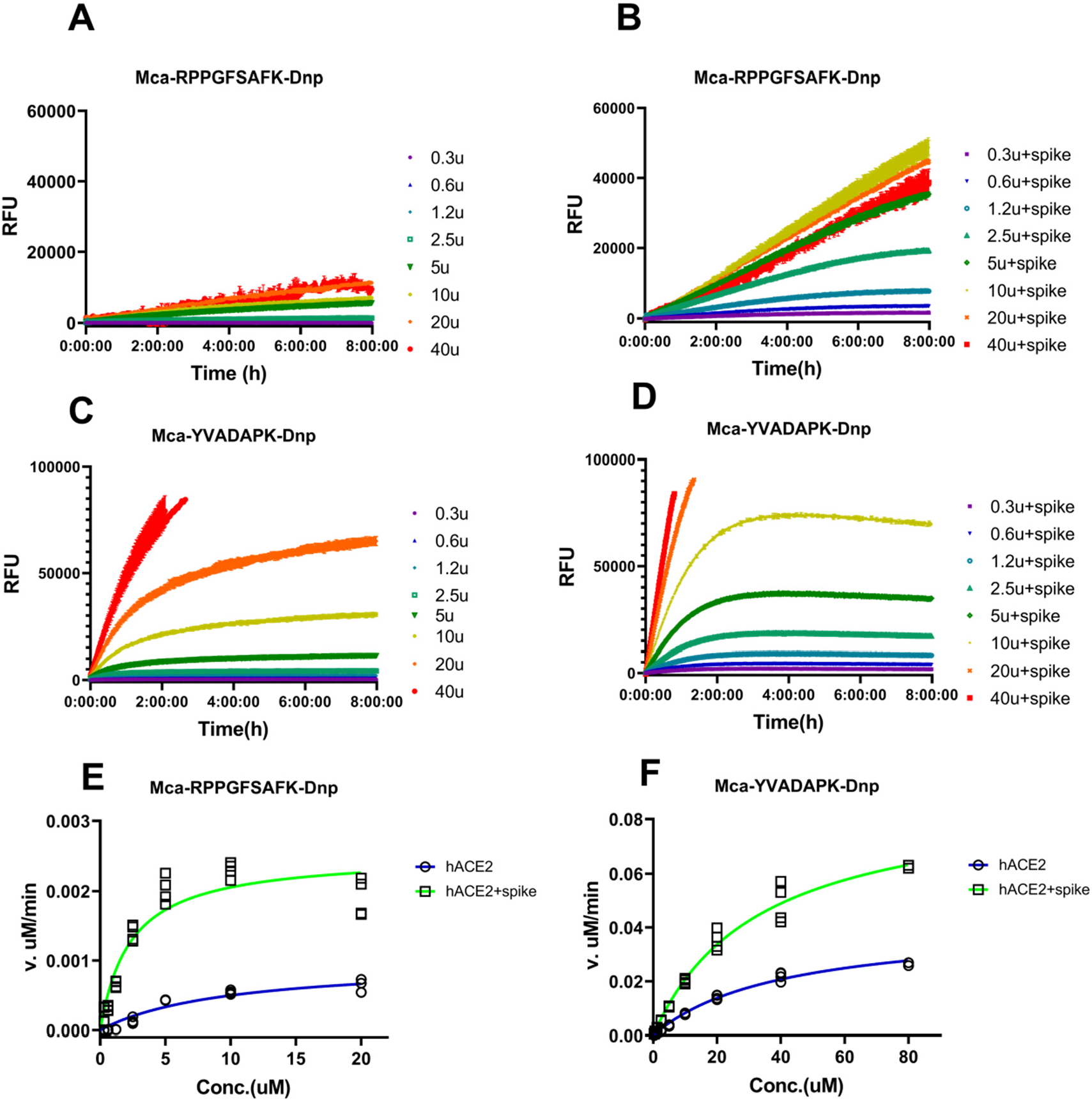
Measurement of kinetic constants for hydrolysis of Mca-RPPGFSAFK-Dnp and Mca-YVADAPK-Dnp in the absence or presence of SARS-CoV-2 spike protein (14ug/ml). A) and B) background subtracted RFU kinetic readings during ACE2 hydrolysis of Mca-RPPGFSAFK-Dnp at different concentrations in the absence (A) and presence (B) of SARS-CoV-2 spike protein. C) and D) background subtracted RFU kinetic readings during ACE2 hydrolysis of Mca-YVADAPK-Dnp at different concentrations in the absence (C) and presence (D) of SARS-CoV-2 spike protein. E) and F) Michaelis plots for ACE2 hydrolysis of Mca-RPPGFSAFK-Dnp and Mca-YVADAPK-Dnp in the absence (blue) or presence (green)of SARS-CoV-2 spike protein. The initial velocity conditions were limited to 30min for Mca-YVADAPK-Dnp and 60min for Mca-RPPGFSAFK-Dnp due to different cleavage rate. All determinations were repeated with duplicates.

**Table 1.** Kinetics constants for hydrolysis of fluorogenic peptides Mca-YVADAPK-Dnp and Mca-RPPGFSAFK-Dnp by ACE2 in the presence of SARS-CoV-2 spike protein (14 ug/ml)

| Substrate | *SARS-CoV-2 Spike*<br>(14ug/ml) | *NaCl (mM)* | $K_m$<br>(uM) | $k_{cat}$<br>(s<sup>-1</sup>) | $k_{cat}/K_m$<br>(M<sup>-1</sup>s<sup>-1</sup>) | $K_i$ (uM) for Competitive substrates | | |
| --- | --- | --- | --- | --- | --- | --- | --- | --- |
|  |  |  |  |  |  | *BK* | *desBK* | *Ang II* |
| YVADAPK | - | 150 | 46.6±10.3 | 0.33±0.05 | 7.08×10<sup>3</sup> | n.d | n.d | 35 ±3 |
|  | + | 150 | 28.2±0.6 | 0.58±0.07 | 2.06×10<sup>4</sup> | 457 ±15 | 1180 ±605 | 5.2± 0.1 |
|  | - | 300 | 24.8±9.9 | 0.058±0.01 | 1.66×10<sup>4</sup> |  |  |  |
|  | + | 300 | 21.1±6.8 | 0.11±0.014 | 3.71×10<sup>4</sup> |  |  |  |
| RPPGFSAFK | - | 150 | 10.2±2.7 | 0.007±0.002 | 7.06×10<sup>2</sup> | n.d | n.d | 57±17 |
|  | + | 150 | 2.41±0.85 | 0.018±0.002 | 7.55×10<sup>3</sup> | n.d | n.d | 4.5 ±1.0 |
|  | - | 300 | 10.0±3.0 | 0.01±0.005 | 1.02×10<sup>3</sup> |  |  |  |
|  | + | 300 | 1.3±0.55 | 0.015±0.004 | 1.14×10<sup>4</sup> |  |  |  |
| Notes: blank cell (not measured); n.d (not detectable) |  |  |  |  |  |  |  |  |

### Competitive ACE2 cleavage in the presence of BK, desBK, and Ang II

From the measurements of kinetic constants of ACE2, we revealed that the binding of SARS-CoV-2 spike protein not only altered the substrate affinity (*K*_m_) to ACE2 but also changed its enzymatic efficiency. In the presence of SARS-CoV-2 spike protein, ACE2 cleaved bradykinin-analog with a *K*_m_ of ~2uM. As previously reported, ACE2 could also cleave des-Arg^9^-bradykinin but with a much higher *K*_m_ (290uM). Therefore, we wonder how the binding of SARS-CoV-2 spike protein would alter the cleavage dynamics among different substrates. We then designed substrate competition assays to assess the selectivity and activity of ACE2 upon binding of SARS-CoV-2 RBD. In the enzymatic reactions, we utilized non-fluorogenic BK, desBK or Ang II peptide to compete for ACE2 cleavage of fluorogenic substrates (**Figure 4**). In the absence of SARS-CoV-2 RBD binding, only Ang II but not BK or desBK inhibited the cleavage of caspase-1 substrate (Mca-YVADAPK(Dnp)-OH) with >95% inhibition at 80uM, which was consistent with the low *K*_m_ (2.0uM) of Ang II and the high *K*_m_ of desBK or very poor activity of BK (**Figure 4 B, D, F and H**). The competitive ACE2 cleavage of the caspase-1 substrate resulted an inhibition constant Ki for Ang II to be 35 μM and 5.2 μM, respectively, in the absence and presence of SARS-CoV-2 RBD, suggesting the SARS-CoV-2 RBD enhanced ACE2 activity against Ang II. Similarly, BK and desBK inhibited the ACE2 cleavage with Ki of 457 μM and 1180 μM, respectively, in the presence of SARS-CoV-2 RBD(**Figure 4 A, C, E and G**), suggesting that SARS-CoV-2 also enhanced ACE2 binding to BK and desBK. As for ACE2 cleavage of the BK-analog, Ang II competed with a Ki constant of 57 μM and 4.5 μM in the absence and presence of the SARS-CoV-2 RBD, respectively (**supplement figure 3**), similar to that from the caspase-1 substrate cleavage. However, BK and desBK failed to inhibit ACE2 cleavage of BK-analog. As shown before, BK-analog bound to endothelin converting enzyme-1 (ECE-1) much better than BK due to its C-terminal modification^16^. It is likely that BK-analog also bound better to ACE2 than BK.

**Figure 4.**
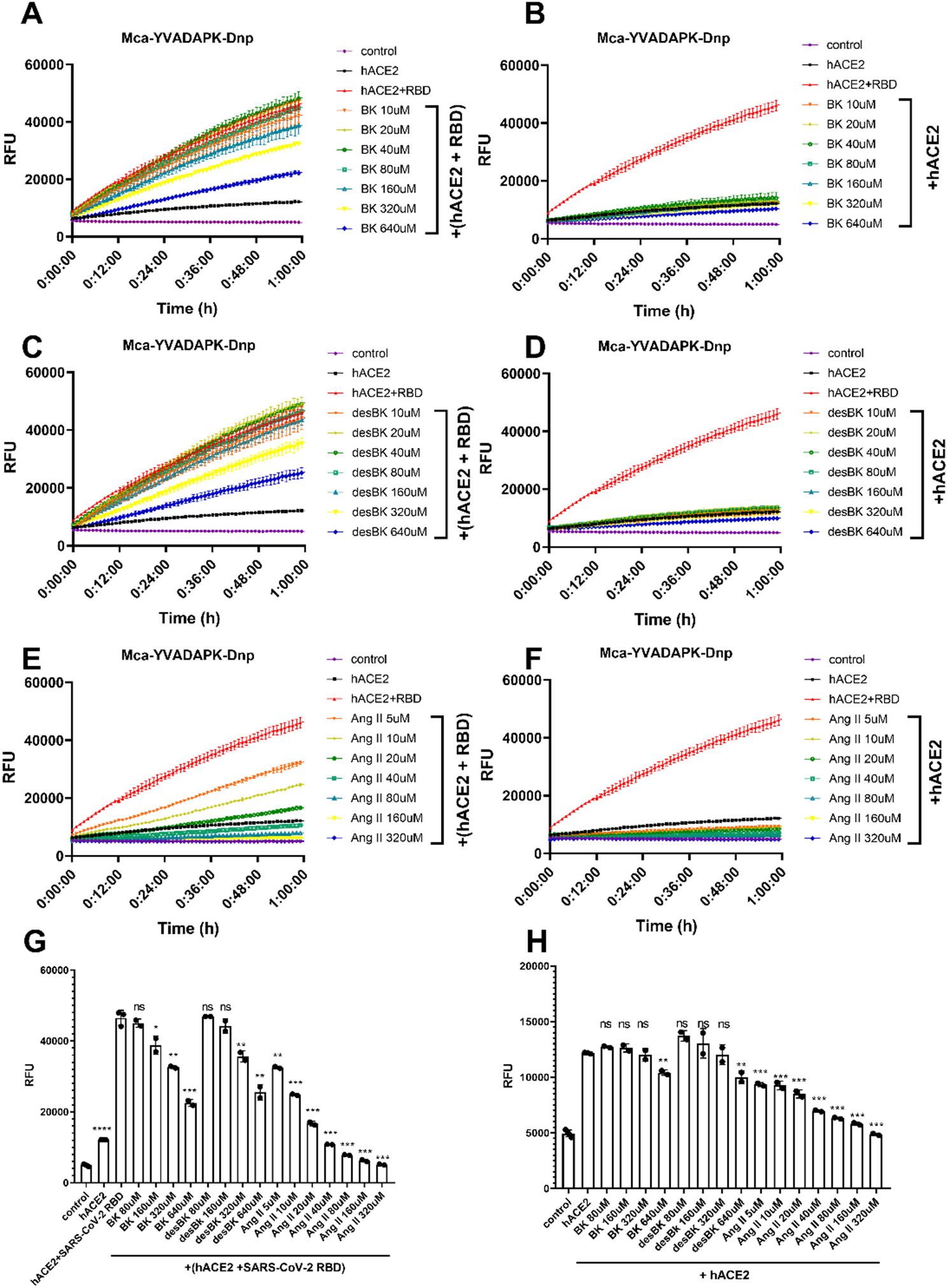
Competition of Bradykinin(BK), des-Arg9-BK, and Ang II peptide with Mca-YVADAPK-Dnp (10uM) for ACE2 hydrolysis. A),C) and E) panels showing the substrates competition in the presence of SARS-CoV-2 RBD protein at different concentrations. B),D) and F) panels showing the substrates competition in the absence of SARS-CoV-2 RBD protein at different concentrations. G) Comparison of RFU during ACE2 hydrolysis of Mca-YVADAPK-Dnp in the presence of competitive substrate at time 1h. The pairwise p-value statistics were calculated between (hACE2 + SARS-CoV-2 RBD) and corresponding concentrations of competitive peptides. H) Comparison of RFU during ACE2 hydrolysis of Mca-YVADAPK-Dnp in the absence of competitive substrate at time 1h. The pairwise p-value statistics were calculated between hACE2 and corresponding concentrations of competitive peptides.

## Discussion

The new coronavirus SARS-CoV-2 infects human cells via the binding of its spike (S) glycoprotein to the human angiotensin-converting enzyme-2 (ACE2). Here we examined the influence of the high affinity binding of SARS-CoV-2 spike protein on the enzymatic activity of ACE2. Surprisingly, our enzymatic assay showed that the SARS-CoV-2 spike protein significantly enhanced the enzymatic activity of ACE2 to cleave YVADAPK peptide, suggesting ACE2 upon SARS-CoV-2 binding would likely cleave more efficiently other substrates with proline at P1 position such as Ang II (DRVYIHP↓F), Apelin-13(QRPRLSHKGPMP↓F). Similarly, binding of SARS-CoV-2 spike enhanced ACE2 to cleave bradykinin analog RPPGFSAFK. In addition, bradykinin (RPPGFSPFR) competed with YVADAPK peptide for ACE2 cleavage to a similar extent as des-Arg^9^-bradykinin. Our results showed that SARS-CoV-2 spike protein not only bound to ACE2 as the entry receptor but also hijacked its enzymatic activities.

We then compared various structures of ACE2 to reveal what structural changes on ACE2 proteolytic domain might happen upon SARS-CoV-2 spike binding to affect the enzymatic activity of ACE2. The native structure of ACE2 extracellular proteolytic domain comprised two subdomains, the N terminal and C terminal subdomain with a wide cleft in between for substrate binding and catalysis^19^, indicating different peptides could be accommodated in deep substrate binding groove (**Figure 5A**). The structure of ACE2 with a potent inhibitor MLN-4760 bound at the active site showed a clear ligand-induced hinge bending movement between the N- and C-terminal domains (**Figure 5B**). Actually this closed conformation of ACE2 proteolytic domain upon inhibitor binding was consistent with ACE2 homolog ACE when one hydrolyzed product Ang II bound to ACE active site^20^, suggesting a common mode to bind substrate and reaction intermediates among zinc metalloproteases (**Figure 5C**). The comparison between native and inhibitor bound ACE2 structures revealed a critical substrate binding pocket including residues F274, H345, F504, H505, Y510 and Y515. Structural studies demonstrated that both SARS-CoV-2 and SARS-CoV spike protein utilized the receptor binding domain (RBD) to recognize ACE2 N-terminal domain^10, 11, 12, 21, 22^. Then we superimposed various SARS-CoV-2 or SARS-CoV RBD bound ACE2 structures onto native ACE2 structure. Structural alignment revealed that both SARS-CoV-2 and SARS-CoV RBD binding did not result in significant conformational changes on the N-domain of ACE2 with r.m.s.d less than 0.5Å (**Figure 5 D and E**). However, the SARS-CoV-2 RBD but not SARS-CoV RBD binding caused the hinge movement between the N- and C-terminal domains reminiscent of a closing clam. We then assessed such relative movement by measuring the angle formed by Asn136 on the rim of native ACE2 C-terminal domain, zinc atom at the catalytic center, and Asn136 of superimposed ACE2 bound by RBD of SARS-CoV-2 or SARS-CoV. Compared to SARS-CoV RBD binding (0.3°), this angle upon SARS-CoV-2 RBD binding was increased to 5°. Particularly, the binding of a chimeric SARS-CoV-2 RBD to ACE2 ^11^, could cause significant movement up to ~12°of C-terminal subdomain toward the N-terminal domain of ACE2 (**Figure 5F**). Furthermore, this angle caused by inhibitor MLN-4760 binding was 29° to form a closed conformation of ACE2, suggesting that SARS-CoV-2 spike binding initiated a progressive conformational change to energetically facilitate ACE2 catalysis. The comparison of substrate binding pocket on ACE2 showed that residues F274, H345, F504, H505, Y510 and Y515 in SARS-CoV-2-ACE2 complexes moved closer to aligned Ang II or MLN-4760 inhibitor, suggesting that SARS-CoV-2 RBD binding to ACE2 could increase the substrate binding (**Figure 5G**). This is consistent with our kinetic constant measurement, which revealed that the *K*_m_ values of ACE2 hydrolysis were decrease upon SARS-CoV-2 spike binding. In addition, in SARS-CoV-2 RBD bound ACE2, H345 residue that was proposed to stabilize the hybridized nitrogen upon catalysis adopted the same conformation as that bound by inhibitor. This active conformation of H345 was not observed in native or SARS-CoV RBD bound ACE2 structure, suggesting that the tighter binding of SARS-CoV-2 RBD could also induce a conformation change to position ACE2 substrate pocket residues for efficient catalysis.

**Figure 5.**
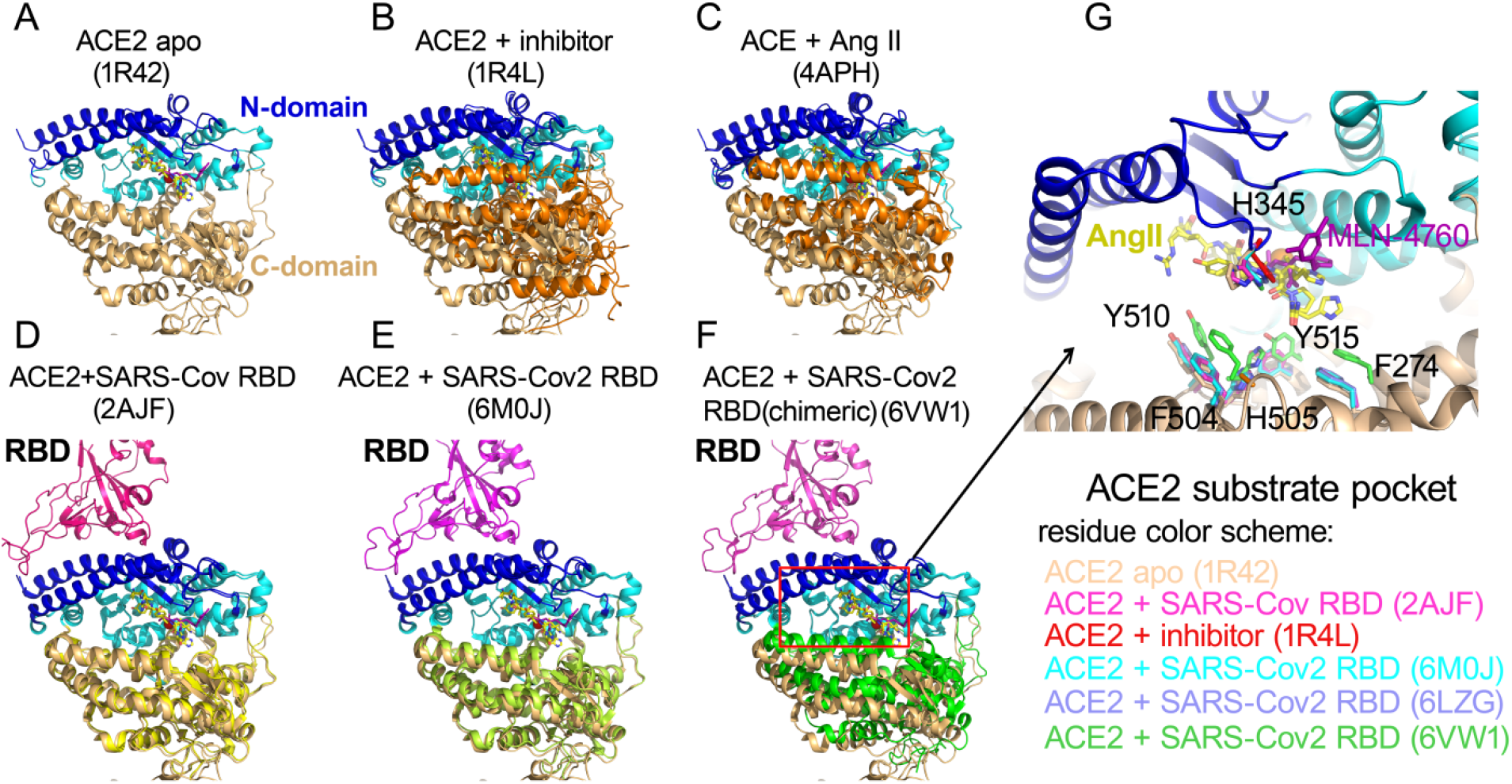
Conformational changes of ACE2 upon SARS-CoV-2 binding. A) apo structure of ACE2 (PDB ID: 1R42).The N-terminal subdomain was colored in cyan with the secondary structures of SARS-CoV-2 spike binding sites highlighted in blue. The C-terminal domain of apo ACE2 was colored in wheat. All subsequent structural superpositions were based the alignment of ACE2 residues 20-84 that formed the first 2 helix for RBD domain interaction. The ACE2 inhibitor MLN-4760 (purple) and Angiotensin II (yellow) were positioned in the substrate pocket based on the structural alignment. B) ACE2 structure with inhibitor MLN-4760 binding (PDB ID: 1R4L). The C-terminal domain was highlighted in orange. C) hACE in complex with Ang II(PDB ID: 4APH). The C-terminal domain was highlighted in orange. D)ACE2 structure in complex with SARS-CoV RBD(PDB ID: 2AJF). The C-terminal domain was highlighted in yellow. E) ACE2 structure in complex with SARS-CoV-2 RBD (PDB ID: 6M0J). The C-terminal domain was highlighted in lime green. F) ACE2 structure in complex with a chimeric SARS-CoV-2 RBD domain (PDB ID: 6VW1). The C-terminal domain was highlighted in green. G) Enlarged view of ACE2 substrate binding pocket. One additional ACE2-SARS-CoV-2 RBD complex(PDB ID: 6LZG) was included. The residue color scheme was listed in the bottom panel.

With the increasing number of confirmed cases worldwide, the major manifestation of SARS-CoV-2 mediated COVID-19 disease and its complications generated critical concerns due to the lack of highly effective antiviral drugs or vaccines. Our current analysis of ACE2 enzymatic activity upon SARS-CoV-2 spike protein binding would be clinically relevant. However, a limitation of our study is that we could not systemically measure the levels of various ACE2 substrates and their cleaved products in COVID-19 patients currently, but we would propose that the ratios between vasodilators and vasoconstrictors would change during SARS-CoV-2 infection due to the high affinity interaction between ACE2 and SARS-CoV-2 spike protein and subsequently altered ACE2 enzymatic activity and substrate selectivity.

Compared to previous outbreaks of SARS-CoV and MERS, SARS-CoV-2 is more contagious and clinical symptoms of COVID-19 are more variable^4^. In addition to the accumulation of fluid in the lung, a considerable fraction of critically ill COVID-19 patients developed deep venous thrombosis or pulmonary embolism^5^. These clinical observations share a common prognostic feature that pointed to the cardiovascular system. Vasodilation, vasoconstriction, and permeability are the essential components of the cardiovascular system, which are regulated by the crosstalk of many hormones and biological active peptides. Among them, renin-angiotensin system and kinin-kellikrein system have major effects upon the cardiovascular systems^14, 17^. Importantly, ACE2 catalyzes the conversion of these regulatory peptides in these systems to maintain the homeostatic state. Among the biological active substrates, ACE2 cleaves Ang II (DRVYIHP↓F), Apelin-13(QRPRLSHKGPMP↓F), and dynorphin A 1-13(YGGFLRRIRPKL↓K) with high proteolytic efficiency (K_m_ <10uM and *k*_cat_/*K*_m_ >1×10^6^M^−1^s^−1^). ACE2 also has substantial activity on Ang I (DRVYIHPFH↓L) des-Arg^9^-bradykinin (RPPGFSP↓F), caspase-1 substrate (Mca-YVADAP↓K) and neurotensin 1-8 (ELYENKP↓R) but with much high *K*_m_ values(~100-300uM). ACE2 hydrolyzes Ang I (DRVYIHPFH↓L) very slowly and does not hydrolyze Bradykinin (RPPGFSPFR) under physiological condition. Characterization of the enzymatic activity of ACE2 demonstrated that SARS-CoV-2 significantly increased substrate affinities by lowering *K*_m_ values and also boosted efficiency (*k*_cat_/*K*_m_) on different substrates. The alteration of kinetic constants of ACE2 enzymatic activity may change the pharmacological dynamics of the substrate and product of ACE2-catalyzed hydrolysis *in vivo*.

Among the peptide components of the rennin-angiotensin system, ACE2 could efficiently hydrolyze Ang II (DRVYIHP↓F), and partially cleave Ang I(DRVYIHPFH↓L). During SARS-CoV-2 infection, the enhancement of ACE2 activity could produce more of their cleaved products, Ang 1-9 and Ang 1-7. Beside the opposing effects of Ang II and Ang 1-7 on vasoconstriction and vasodilatation, they regulated the expression of ACE2. Several *in vivo* and *in vitro* studies documented that Ang II decreased the expression of ACE2 through activation of MAPK pathways, whereas Ang 1-7 counteracted the inhibitory effect of Ang II to boost the expression of ACE2 through MAPK phosphatase activation^23, 24^. Previous studies showed that SARS-CoV infection caused the internalization of ACE2^6^. However, analyzing the lung tissue in ACE2 mice 3 days post SARS-CoV-2 infection showed substantial expression of ACE2^7^. More importantly, ACE2 expression was increased in the lungs of severe COVID-19 patients with comorbidities, compared to control individuals^18, 25^. The enhanced ACE2 activity to cleave Ang II upon SARS-CoV-2 spike binding might be one of the mechanisms whereby SARS-CoV-2 utilized to boost the expression of ACE2.

Apelin 1-13(QRPRLSHKGPMP↓F) is the endogenous ligand for the angiotensin II protein J receptor (APJR), which is a homolog of the angiotensin receptor AT1^26^. Apelin 1-13 has nanomolar affinity to APJR and the major effect of Apelin *in vivo* was nitric oxide dependent arterial dilatation to counterbalance Ang II effects. The cleavage of its carboxyl-terminal phenylalanine of Apelin 1-13 cause the loss of critical interactions between Phe13 and a large hydrophobic cavity on APJR^27^, resulting in 2-5 fold decrease of its potency^28^. The enhancement of ACE2 activity would accelerate the metabolism of Apelin 1-13, further dampening its beneficial effects on overall cardiac functions.

The proteolysis of bradykinin may be particularly relevant to COVID-19. Normally ACE is responsible for the hydrolysis of bradykinin, whereas ACE2 hydrolyzes des-Arg^9^-bradykinin^14^. Bradykinin selectively binds to B2 receptors and des-Arg^9^-bradykinin selectively binds to B1 receptor. Compared to the constitutive expression of B2 receptor, expression of B1 receptor is upregulated during inflammation. In addition, previous studies showed that all of the reactions for bradykinin degradation occurred normally but the production rate of des-Arg^9^-bradykinin was accelerated fivefold when blood was clotted^29^. Our results showed that SARS-CoV-2 spike protein enhanced bradykinin and des-Arg^9^-bradykinin binding to ACE2. Furthermore, neuropeptides Dynorphin A 1-13(YGGFLRRIRPKL↓K) and β– Casomorphin(YPFVEP↓I) are good ACE2 substrate, which are endogenous ligands for opioid receptors and have antinociceptive effects^30^. Interestingly, Dynorphin A 1-13 was also implicated as an appetite stimulant^31^. The removal of the C-terminal lysine residue of Dynorphin A 1-13 resulted in 90% loss of its potency^32^. Since symptoms of COVID-19 also include body aches and loss of taste, it would be intriguing to speculate that altered ACE2 hydrolysis of these neuropeptides would affect the homeostatic functions of these opioid receptors as well.

Collectively, the homeostatic regulation of cardiovascular system controlled by ACE2 would be out of balance due to the altered activity upon SARS-CoV-2 spike protein binding, resulting in further complications of SARS-CoV-2 viral infection. Our study documented the direct influence of SARS-CoV-2 infection on the enzymatic activity of ACE2 and merited the further investigation of biological roles of ACE2 upon this new coronavirus infection, which may shed new light on the pathogenesis of COVID-19 and development of better treatment of this disease and its complications.

## Methods

### Reagents

Recombinant proteins of human ACE2 extracellular enzymatic domain were purchased from R&D systems(933-ZN) and Sinobiological, Inc(10108-H08H). Recombinant proteins of SARS-CoV RBD (40150-V08B2) and SARS-CoV-2 RBD proteins (40592-V08H) were obtained from Sinobiological, Inc. Fluorogenic peptide substrates were obtained from R&D systems, where Mca-YVADAPK(Dnp)-OH (ES007) was used as an angiotensin II like substrate and Mca-RPPGFSAFK(Dnp)-OH(ES005) was a fluorogenic peptide derivative of Bradykinin according to manufacturer’s instruction. Non-fluorogenic bradykinin(BK, RPPGFSPFR), des-Arg^9^-bradykinin (desBK, RPPGFSPF) and angiotensin II (Ang II, DRVYIHPF) peptides were purchased from Genscript, Inc. Fluorogenic peptide Mca-Pro-Leu-OH (Bachem, M-1975) was used as calibration standard to obtain the molarity (pmol) to relative fluorescence unit (RFU) conversion factor on Synergy H1 fluorescent plate reader.

### SARS-CoV-2 trimeric spike protein expression and purification

The vector to express the prefusion S ectodomain, a gene encoding residues 1-1208 of SARS-CoV-2(GenBank: MN908947) was kindly provided by Jason S. McLellan (the University of Texas at Austin) and the protein was expressed as described previously^9^. In brief, FreeStyle293 F cells (Thermo Fisher) were transiently transfected with this expression vector using polyethylenimine (PEI). Protein was purified from filtered cell supernatants using StrepTactin XT resin (IBA, Inc) and subsequently subjected to further purification by size exclusion chromatography using a Superose 6 10/300 column (GE Healthcare) with the buffer of 75mM Tris (pH7.5), 0.15M NaCl.

Before enzymatic assays, SARS-CoV-2 spike and commercial RBD recombinant proteins were dialyzed extensively against 75mM Tris(pH7.5) and 0.15M NaCl.

### Enzymatic assays of ACE2 hydrolysis of biological peptides

Reactions were performed in black microtiter plates at ambient temperature (26°C) according to manufacturer’s instruction. To each well, 50ul of 0.2 or 0.4ug/ml ACE2 in assay buffer containing 75mM Tris(pH7.5) plus NaCl at 0.15M, 0.3M or 1.0M were added respectively. Then 10ul dialysis buffer or SARS-CoV-2 S spike/RBD domain proteins at various final concentrations ranging from 70ug/ml to 0.4ug/ml were added to wells and incubated for 20min after mixing by shaking. The reactions were initiated by adding 50ul of fluorogenic peptides at 40uM or with 2-fold serial dilutions ranging from 160uM to 0.6uM to determine the kinetic constants for ACE2 hydrolysis. The relative fluorescence units (RFU) were read at excitation and emission wavelengths of 320nm and 405nm (top read), respectively in kinetic mode at 1-minute intervals for 8 hours.

To calculate specific activity of ACE2, the substrate blank adjusted relative fluorescent units were converted to molarities according to the conversion factor on Synergy H1 plate reader that was calibrated by standard Mca-pro-Leu-OH (**supplement figure 2**). To obtain the kinetic constants, the hydrolysis was limited to ≤15% product formed as the initial velocity conditions. Initial velocities were plotted versus substrate concentration and fit to the Mechaelis-Menten equation *v*=Vmax[S]/(*K*_m_ +[S]) using GraphPad Prism software. Turnover numbers (*k*_cat_) were calculated from the equation *k*_cat_=Vmax/[E], using a calculated ACE2 molecular mass of 85,314 Da and assuming the enzyme sample to be essentially pure and fully active.

### Substrate competition assays using non-fluorogenic BK, desBK and Ang II peptides

To compare the relative proteolytic activity of ACE2 to different substrates, various concentrations of non-fluorogenic BK, desBK or Ang II peptides were added to the reaction mixture with fluorogenic peptides as competitive substrates. As mentioned above, 50ul of ACE2 at 0.4ug/ml was preincubated with buffers or SARS-CoV-2 RBD protein at a final concentration of 125nM for 20min. Subsequently, 50ul of fluorogenic peptides at 20uM supplemented with a serial dilution of non-fluorogenic BK, desBK or Ang II peptides ranging 640uM to 80uM were added to initiate enzymatic cleavage. Fluorescence changes were measured at 1min intervals for 1 hour at 26 degree as the initial velocity conditions.

### Statistical analysis

Student’s t-Test was used to determine the statistical significance in each pair wise comparison. Each experiment was repeated at least with duplicates. P<0.05 was considered statistically significant(* P<0.05; ** P<0.01; ***P<0.001; **** P<0.0001).

## Disclosure

The authors declare that no competing interests exist.

## Author contributions

J. Lu conceived this project and carried out all the experiments. J. Lu and P. Sun wrote the manuscript.

## Acknowledgement

The funding of this work is provided by the Intramural Research Program (IRP) of National Institute of Allergy and Infectious Diseases (NIAID), National Institutes of Health.

---

**Supplementary figure 1.**
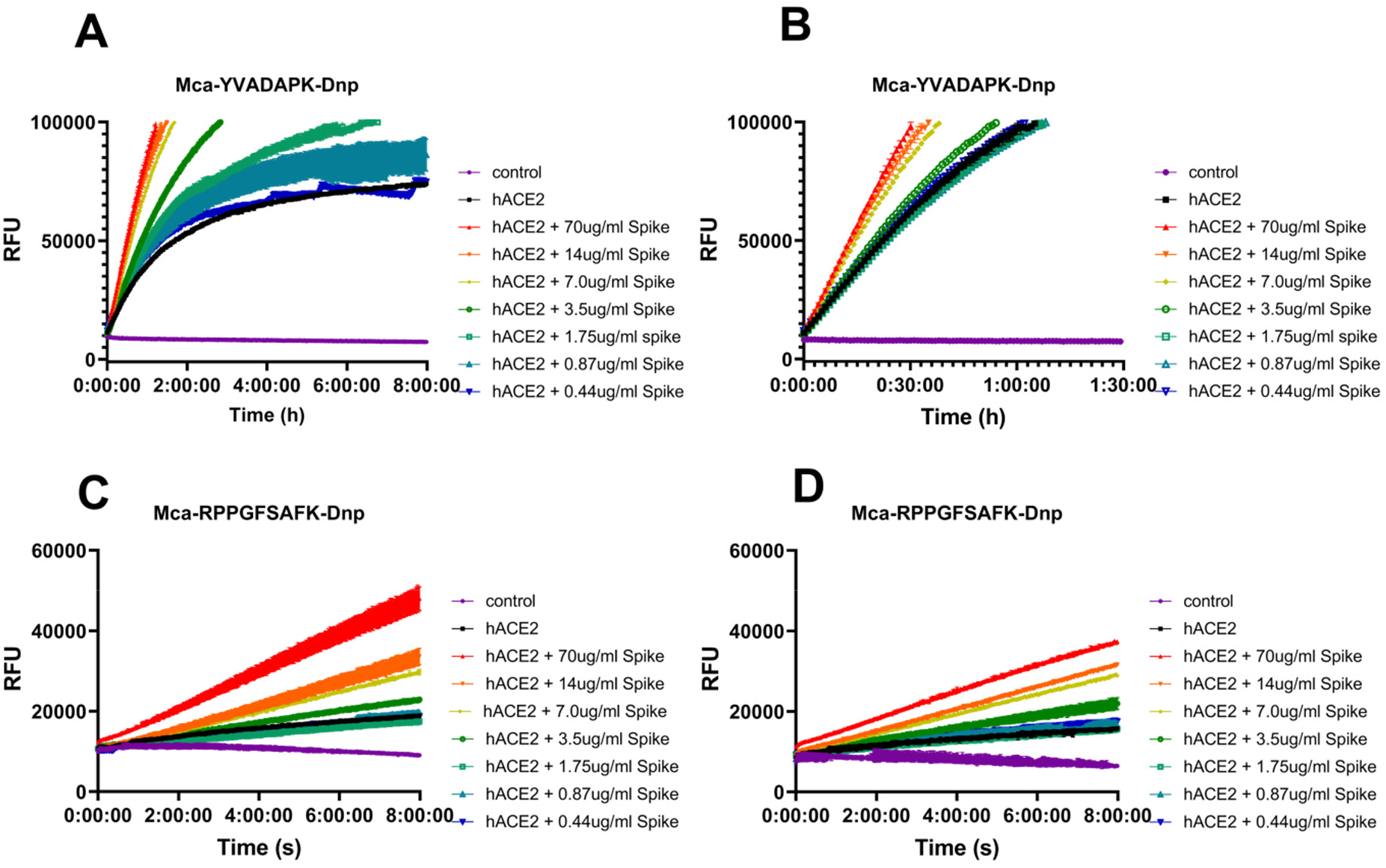
SARS-CoV-2 spike protein enhanced ACE2 cleavage of fluorogenic caspase-1 substrate and bradykinin analog in a concentration dependent manner with increased concentration of NaCl in the assay buffer. A) and C) Kinetic reading of relative fluorescence units (RFU) monitoring the hydrolysis of Mca-YVADAPK-Dnp(20uM) and Mca-RPPGFSAFK-Dnp (20uM) in the presence of SARS-CoV-2 spike protein at indicated final concentrations in the enzymatic assays with 0.3M NaCl. The maximum reading is limited to 100000 RFU on Synergy H1 plate reader. B) and D) Kinetic reading of relative fluorescence units (RFU) monitoring the hydrolysis of Mca-YVADAPK-Dnp(20uM) and Mca-RPPGFSAFK-Dnp (20uM) in the presence of SARS-CoV-2 spike protein at indicated final concentrations in the enzymatic assays with 1.0M NaCl.

**Supplementary figure 2.**
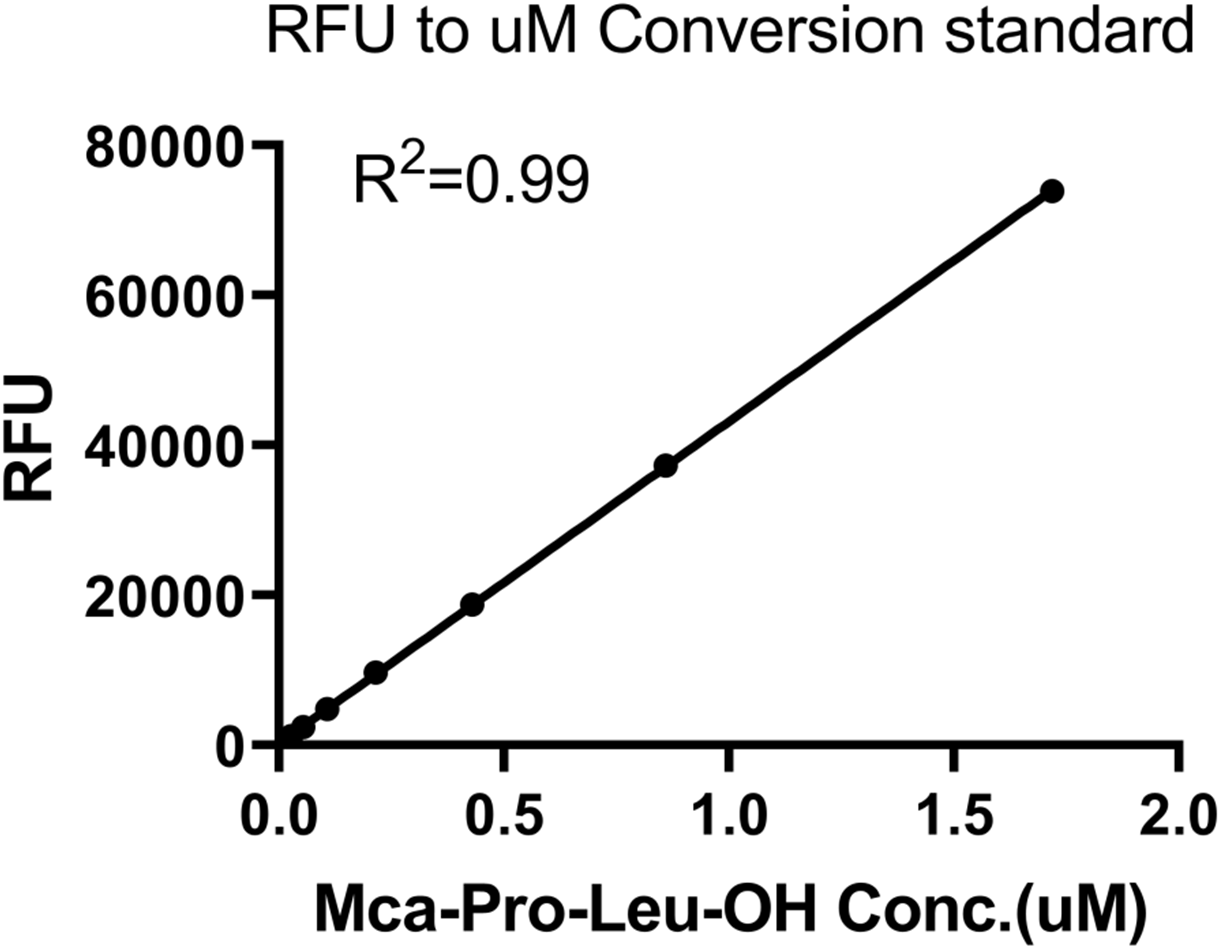
Calibration of Synergy H1 plate reader with fluorescent Mca-Pro-Leu-OH peptide to obtain relative fluorescent unit to molarity conversion factor. All readings were carried out in 110 ul of enzymatic assay buffers with various concentrations of Mca-Pro-Leu-OH peptide.

**Supplementary figure 3.**
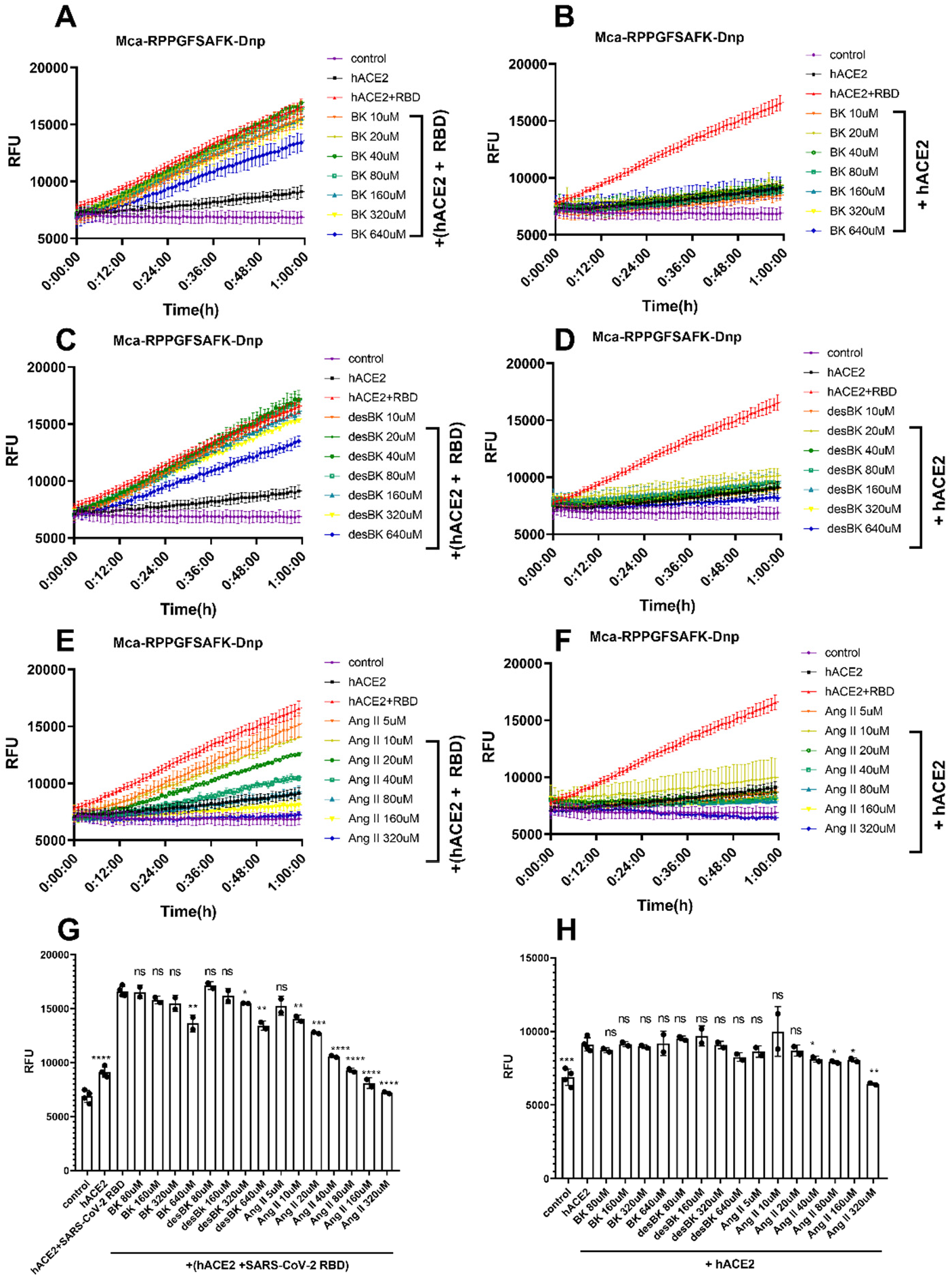
Competition of Bradykinin(BK), des-Arg9-BK, and Ang II peptide with Mca-RPPGFSAFK-Dnp (10uM) for ACE2 hydrolysis. A),C) and E) panels showing the substrates competition in the presence of SARS-CoV-2 RBD protein at different concentrations. B),D) and F) panels showing the substrates competition in the absence of SARS-CoV-2 RBD protein at different concentrations. G) Comparison of RFU during ACE2 hydrolysis of Mca-RPPGFSAFK-Dnp in the presence of competitive substrate at time 1h. The pairwise p-value statistics were calculated between (hACE2 + SARS-CoV-2 RBD) and corresponding concentrations of competitive peptides. H) Comparison of RFU during ACE2 hydrolysis of Mca-RPPGFSAFK-Dnp in the absence of competitive substrate at time 1h. The pairwise p-value statistics were calculated between hACE2 and corresponding concentrations of competitive peptides.

## Notes

### Competing Interest Statement

The authors have declared no competing interest.

